# Inhibition of microglial GBA hampers the microglia-mediated anti-oxidant and protective response in neurons

**DOI:** 10.1101/2021.01.20.427380

**Authors:** Electra Brunialti, Alessandro Villa, Marianna Mekhaeil, Federica Mornata, Elisabetta Vegeto, Adriana Maggi, Donato A. Di Monte, Paolo Ciana

## Abstract

Homozygotic mutations in the GBA gene cause Gaucher’s disease, moreover, both patients and heterozygotic carriers have been associated with 20- to 30-fold increased risk of developing Parkinson’s disease. In homozygosis, these mutations impair the activity of β-glucocerebrosidase, the enzyme encoded by GBA, and generate a lysosomal disorder in macrophages, which changes morphology towards an engorged phenotype, considered the hallmark of Gaucher’s disease. In the brain, most of the pathological effects caused by GBA mutations have been attributed to the β-glucocerebrosidase deficit in neurons, while a microglial phenotype for these mutations has never been reported. Here, we applied the bioluminescence imaging technology, immunohistochemical and gene expression analysis to investigate the consequences of microglial β-glucocerebrosidase inhibition in the brain of reporter mice, in primary neuron/microglia co-cultures and in cell lines. Our data demonstrate the existence of a novel mechanism by which microglia sustain the antioxidant/detoxifying response mediated by the nuclear factor erythroid 2-related factor 2 in neurons. The central role played by microglia in this neuronal response *in vivo* was proven by pharmacological depletion of the lineage in the brain, while co-cultures experiments provided insight on the nature of this cell-to-cell communication showing that this mechanism requires a direct microglia-to-neuron contact supported by functional actin structures. Pharmacological inhibition of microglial β-glucocerebrosidase was proven to induce morphological changes, turn on an anti-inflammatory/repairing pathway and hinder the microglia ability to activate the anti-oxidant/detoxifying response, thus increasing the neuronal susceptibility to neurotoxins.

Altogether, our data suggest that microglial β-glucocerebrosidase inhibition impairs microglia-to-neuron communication increasing the sensitivity of neurons to oxidative or toxic insults, thus providing a possible mechanism for the increased risk of neurodegeneration observed in carriers of GBA mutations.

**Graphical Abstract:** 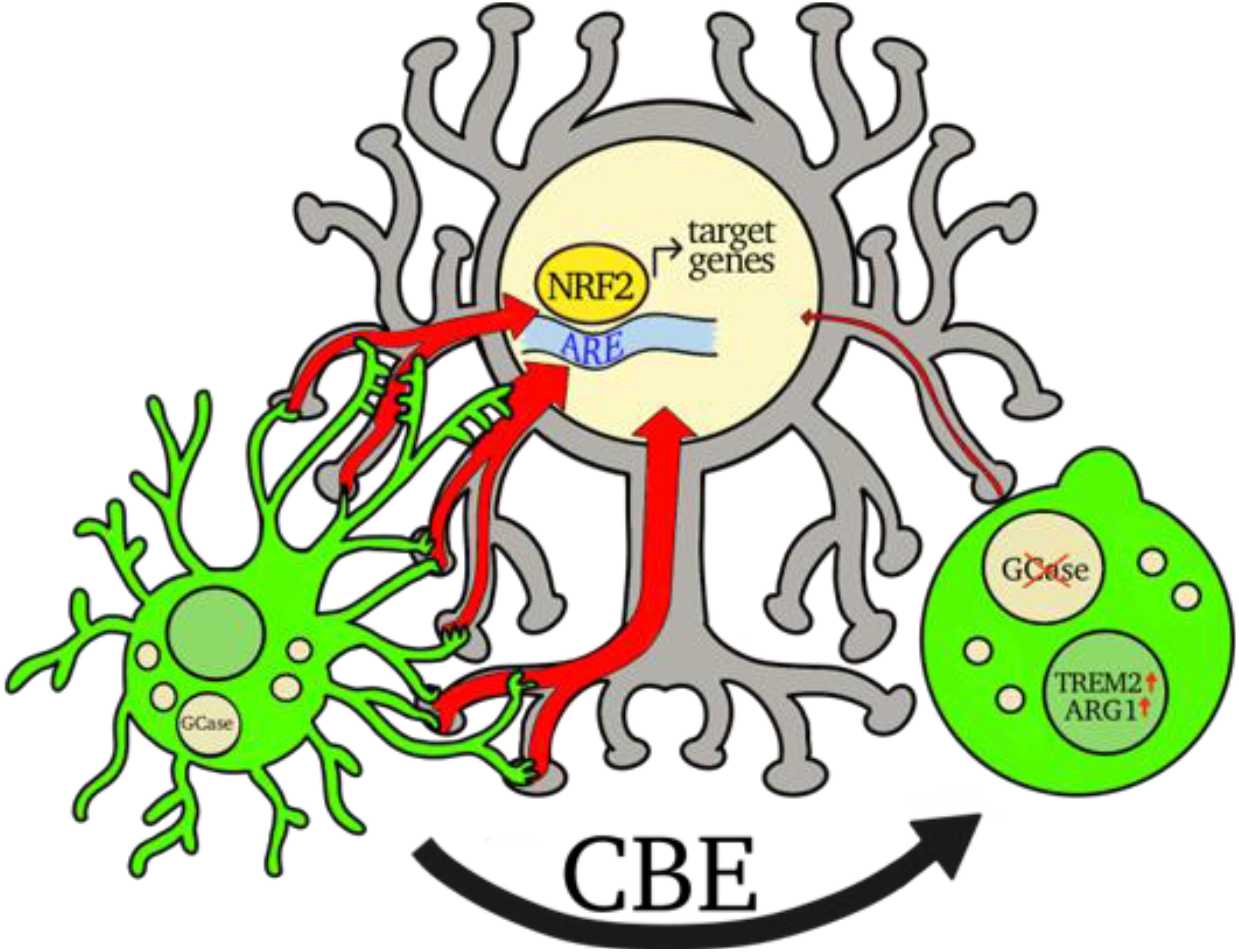

**In Brief:** Microglia, through actin-dependent structures, contact neurons and induce a detoxification response by increasing the NFE2L2 signalling pathway. Inhibition of GCase activity by CBE treatment produces a morpho-functional change in microglia cells hampering the neuroprotective microglia-neuron communication thus inducing a phenotype in dopaminergic neurons characterized by increased susceptibility to oxidative stress or toxic insults.

## Introduction

The human GBA gene is spanning 7.6 Kb of the 1q21 region and consists of 11 exons (Horowitz *et al.*, 1989); approximately 300 pathogenic mutations in the coding regions have been described, each affecting, to a different extent, the GCase activity ranging from negligible to a severe reduction (Hruska *et al.*, 2008). The GCase enzymatic deficiency in homozygous individuals leads to the accumulation of glucosylceramide in lysosomes of the macrophage/monocyte cells causing Gaucher’s disease (Beutler, 1992). In both homozygous and heterozygous carriers, GBA mutations, with different penetrance, display a CNS phenotype that has been associated with neurological symptoms in Gaucher’s disease (Erikson *et al.*, 1997), Lewy body dementia and severe forms of Parkinson’s disease (Sidransky *et al.*, 2009; Bultron *et al.*, 2010; Mata *et al.*, 2016).

GBA mutations represent the most significant genetic risk factor for Parkinson’s disease, since carriers display from 20 to 30 fold increased risk (depending on the genetic variant) of developing the illness, with those variants associated with the most severe neuropathic form of Gaucher’s disease displaying the highest risk (Gan-Or *et al.*, 2015). Several hypotheses have been formulated for the involvement of the GCase impairment in Parkinson’s neurodegeneration, mostly focussing on the role of GBA mutations in neurons.

Although, enzymatic dysfunction leading to the progressive accumulation of glucocerebroside plays a central role in Gaucher’s pathogenesis, substrate accumulation is believed to require mutations in both alleles, thus it is considered unlikely to play a role in Parkinson’s disease associated with GBA mutations (Gegg *et al.*, 2015). Nevertheless, a reciprocal relationship between α-synuclein levels and GCase mutations and/or GCase inhibition have been reported (Migdalska-Richards and Schapira, 2016). This interrelation goes beyond GBA mutations, since low GCase activity has been associated with increased levels of α-synuclein in the brain and in the cerebrospinal fluid of idiopathic Parkinson’s patients (Parnetti *et al.*, 2014). Some study proposed that a reciprocal positive feed-back loop is instigated by the GCase enzyme inhibition due to the accumulation of the GCase substrate in lysosomes that leads to the increased α-synuclein deposition (Yap *et al.*, 2011; Taguchi *et al.*, 2017). Conversely, high levels of α-synuclein in dopaminergic neurons was demonstrated to inhibit the GCase translocation from the endoplasmic reticulum to lysosomes with the subsequent accumulation of the GCase substrates (Mazzulli *et al.*, 2011). GCase inhibition and α-synuclein accumulation may thus potentiate each other contributing to the neurodegenerative process (Balestrino and Schapira, 2018) associated with a neuroinflammatory response characterized by a frank microglia activation (Rocha *et al.*, 2015; Mus *et al.*, 2019).

In addition to the lysosomal impairment, GCase inhibition may also hinder mitochondrial functions in dopaminergic neurons (Cleeter *et al.*, 2013; Osellame *et al.*, 2013) resulting in the production of reactive oxygen species and free radicals; this condition increases susceptibility to exogenous or endogenous stimuli inducing parkinsonism, including 1-methyl-4-phenyl-1,2,3,6-tetrahydropyridine (MPTP), rotenone, and α-synuclein overexpression (Fishbein *et al.*, 2014; Schöndorf *et al.*, 2014; Noelker *et al.*, 2015; Migdalska-Richards *et al.*, 2017; Kim *et al.*, 2018; Yun *et al.*, 2018; Mus *et al.*, 2019). A well-recognized player in the neuron defense from neurotoxic insults and oxidative stress is the nuclear factor erythroid 2-related factor 2 (NFE2L2, also known as NRF2) transcription factor (Zhang *et al.*, 2013). This leucine zipper protein is normally kept in the cytoplasm in an inhibitory complex with the kelch like-ECH-associated protein 1 (KEAP1) and cullin 3 proteins (McMahon *et al.*, 2006); when stimulated NFE2L2 is released from the complex and translocate into the nucleus, where it binds the promoter and regulate the transcription of target genes (Itoh *et al.*, 1997). Noteworthy, NFE2L2 controls the basal and inducible expression of enzymes involved not only in drug detoxification and redox balance, but also in metabolism, autophagy, lysosomal biogenesis, mitochondrial dysfunctions and neuroinflammation (Hayes and Dinkova-Kostova, 2014), and its dysregulation has been associated with neurodegenerative processes. Recently, we demonstrated that NFE2L2 is early activated during neurodegeneration well before the onset of neuronal death (Rizzi *et al.*, 2018). In the present work, we aimed at clarifying whether GBA impairment may interfere with the activation of this neuroprotective pathway thus turning brain into a condition of increased susceptibility to neurotoxic insults. Our results demonstrate the existence of a novel mechanism of neuroprotection based on the physical microglia/neuron interaction that leads to the activation of the NFE2L2 pathway in neurons; inhibition of microglial GCase activity hampers this communication, suggesting an active role of the microglial GBA gene product in the induction of detoxifying signals in neurons. We show that this microglia/neuron communication is not mediated by soluble factors but requires a direct contact between the two cellular components and functional actin structures in the microglial cells.

## Results

### GCase inhibition impairs Nfe2l2 response *in vivo*

Two separate sets of *in vivo* experiments were carried out to investigate potential interactions between GCase activity and the Nfe2l2 pathway. First, changes in Nfe2l2-dependent oxidative stress response caused by GCase inhibition were assessed by *in vivo* and *ex vivo* imaging of luciferase activity in the brain of ARE-*luc2* reporter mice (Fig. 1); in these transgenic animals, the luciferase reporter is expressed under control of the Nfe2l2 transcription factor (Rizzi *et al.*, 2017, 2018). Two groups of 16 reporter mice were treated i.p. with a daily dose of 100 mg/kg/die CBE or vehicle (PBS) for three days, a dose sufficient to decrease GCase activity down to 50% of baseline levels (Supplementary Fig. 2A): the same reduction has been observed in fibroblasts obtained from heterozygous patients carrying severe GBA mutations (Migdalska-Richards and Schapira, 2016). At day 2, half of the mice from both groups were further treated i.p. with two injections (20 and 16 hours prior to the end of the experiment) of 75 mg/kg tert-butylhydroquinone (tBHQ), a well-known Nfe2l2-inducing agent crossing the blood-brain barrier (Silva-Palacios *et al.*, 2017; Rizzi *et al.*, 2018) (Fig. 1A). Bioluminescent emissions were measured before and after tBHQ administration by *in vivo* imaging sessions (Fig. 1A). Quantitative analysis of photon emission from the head area of CBE- and vehicle-treated mice revealed a comparable luciferase activity, indicating that CBE administration did not affect the physiological Nfe2l2 signaling pathway (Fig. 1B, 1C). tBHQ treatment induced a significant (three-fold) increase in luciferase activity in vehicle-treated animals. Remarkably, however, this Nfe2l2 response was impaired in mice pre-treated with CBE. At the end of these experiments, brains were collected and dissected, and *ex vivo* imaging was carried out on whole brains as well as on 2 to 3 mm-thick brain slices (Fig. 1D). In line with the *in vivo* results, quantification of luciferase activity showed that its increase after administration tBHQ was reduced in CBE-treated animals. This effect reached statistical significance in the whole brain (Fig. 1E) and in brain slices I (−53%), II (−20%), and IV (−21%) (Fig. 1F, 1G, 1I).

**Figure 1.**
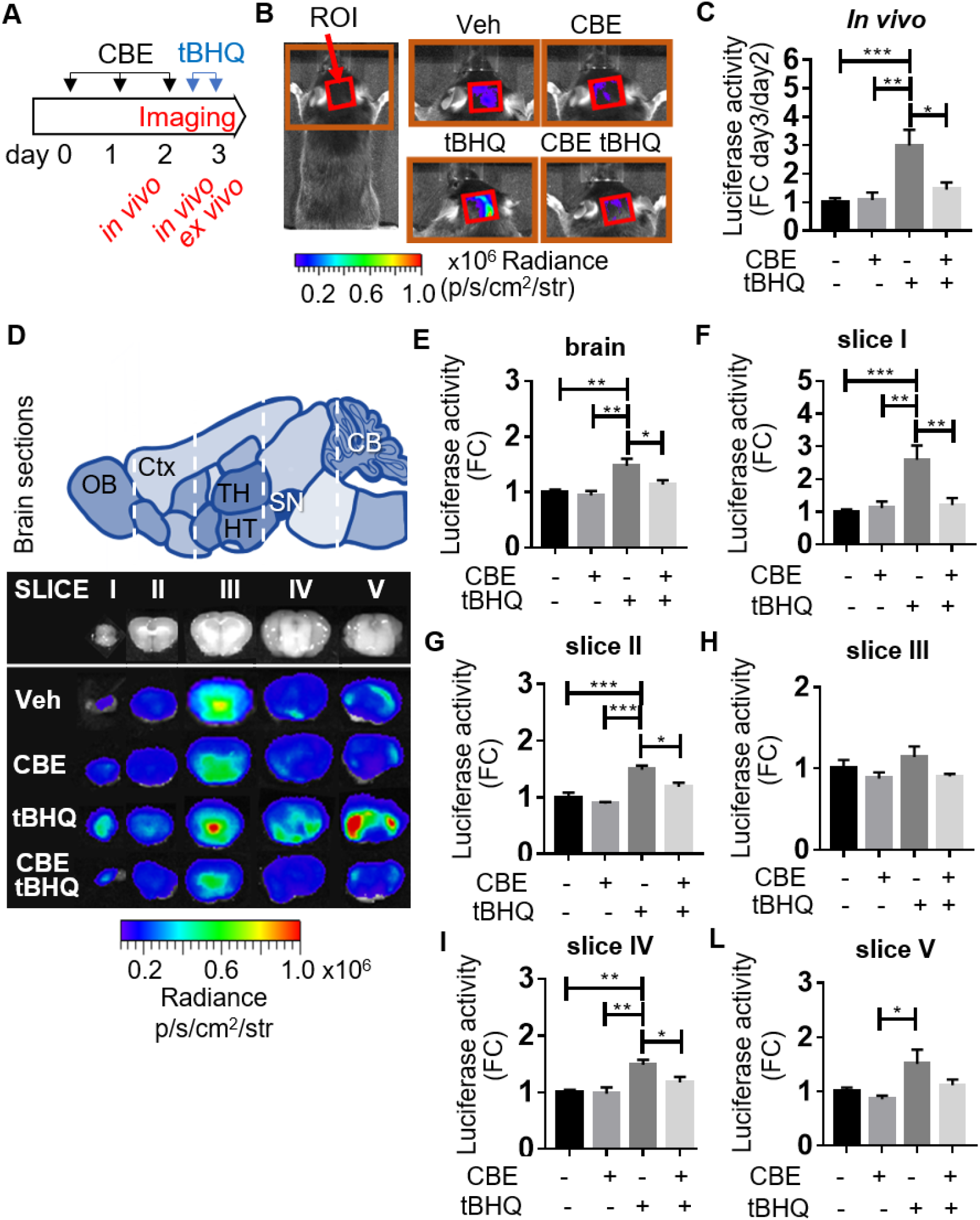
Bioluminescence analysis of Nfe2l2 activation in ARE-*luc2* reporter mice treated with CBE. (**A**) Scheme of the pharmacological treatments and *in vivo/ex vivo* imaging sessions applied to the ARE-*luc2* reporter mouse model. (**B**) Representative pictures of the bioluminescence emitted from the region of interest (ROI) highlighted by the red square on the head area: the intensity of photon emission is reproduced with pseudocolour images generated by the software based on the indicated scale bar. (**C**) The graph reports bioluminescence imaging (BLI) quantifications of the photon emission from the ROI shown in (**B**); the measures of photon emission, radiant flux values (p/s/cm^2^/sr), are expressed in the graph as fold change (FC) *versus* day 2. (**D**) Bioluminescence *ex vivo* imaging analysis of the brain slices (I-V) schematically represented in the upper part together with the different brain areas (CB: cerebellum, Ctx: cerebral cortex, HT: hypothalamus, OB: olfactory bulb, SN: *substantia nigra*, TH: thalamus). In the lower part are shown representative pseudocolur *ex vivo* images of the photon emissions from the indicated brain slices. Quantification of the BLI signals from whole brain and brain slices are reported in (**E**) and (**F-L**) graphs, respectively: radiant flux values (p/s/cm^2^/sr) are expressed as FC *versus* vehicle. Data are mean values ± SEM of n=4 animals measured in duplicate. \**P*<0.05, \*\**P*<0.01, \*\*\**P*<0.001 calculated by one-way ANOVA followed by Tukey’s multiple comparison test.

Next, wild-type C57Bl/6 mice were divided into 4 experimental groups and treated with vehicle, CBE, tBHQ or CBE plus tBHQ as described above. Post-mortem analyses were performed on tissue sections of the ventral mesencephalon that contained the SNpc and were immunostained with anti-tyrosine hydroxylase (Th) and anti-Nfe2l2 antibodies. Immunoreactivity for Nfe2l2 (Red staining) was detected within the cytosolic compartment of Th-positive nigral dopaminergic neurons in sections from vehicle- and CBE-treated mice (Fig. 2A). Quite in contrast, Nfe2l2 staining became mostly nuclear after tBHQ administration, consistent with a nuclear translocation of this transcription factor. Nfe2l2 nuclear immunoreactivity was apparently less robust in the SNpc of animals injected with both CBE and tBHQ, compared to tBHQ alone (Fig. 2A). Semi-quantitative analysis of nuclear Nfe2l2 intensity confirmed these histochemical observations. It showed a 3-fold increase in nuclear Nfe2l2 after treatment with tBHQ alone and a significant reduction (−30%) of this effect in mice with CBE-induced GCase inhibition (Fig. 2B). Taken together, the results of these *in vivo* experiments provided first evidence of a relationship between GCase activity and the Nfe2l2 transcriptional activity, suggesting that impairment of GCase may alter neuronal antioxidant responses in different brain regions including the SNpc. Follow-up studies were then designed to investigate mechanisms underlying this relationship.

**Figure 2:**
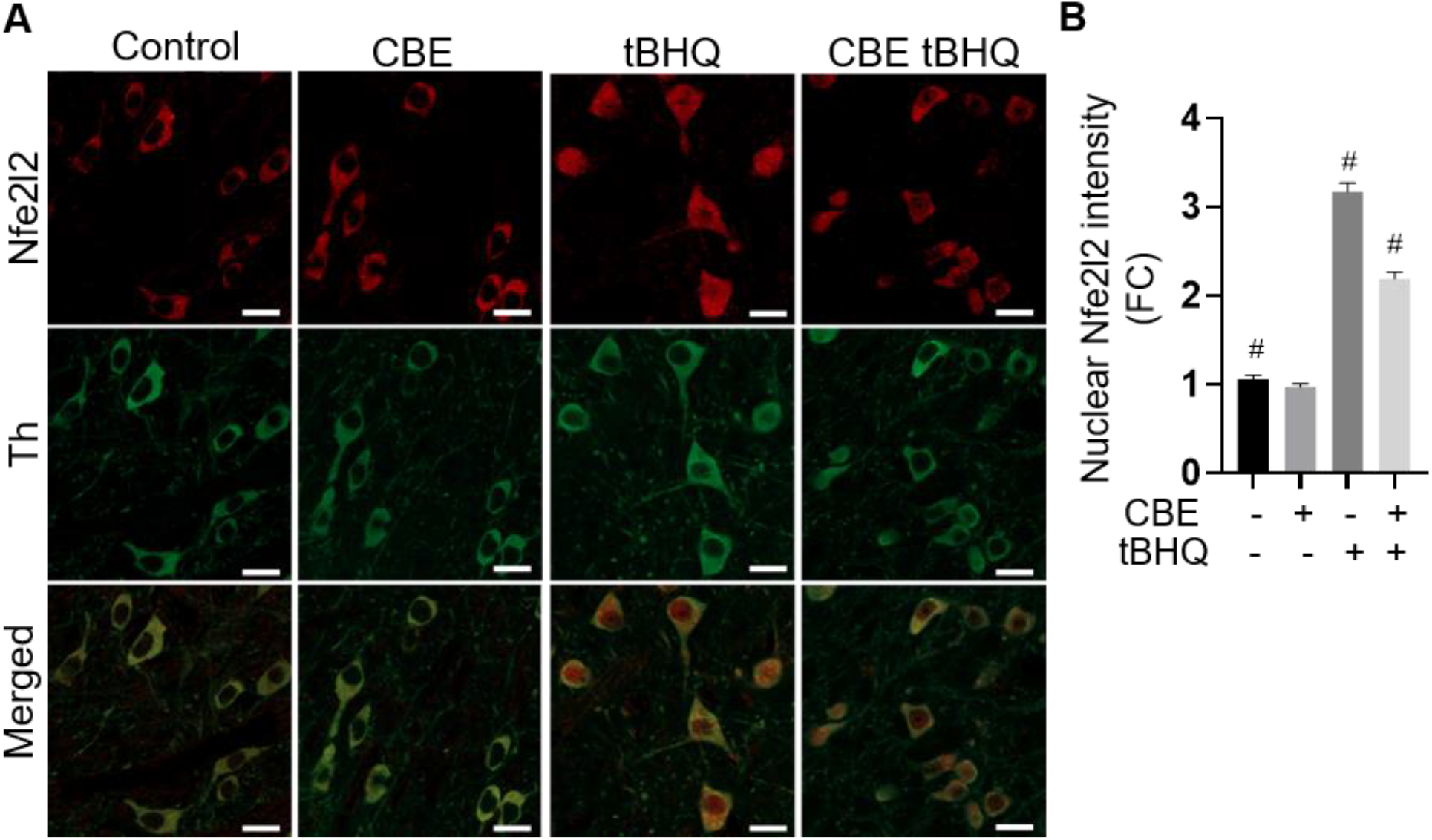
Nfe2l2 activation in nigral dopaminergic neurons. C57BL/6 wild-type mice (n = 4/treatment group) were treated with vehicle (control), CBE, tBHQ or CBE plus tBHQ. Midbrain sections from these animals were double-immunostained with anti-Nfe2l2 and anti-Th. **(A)** Representative confocal images of the substantia nigra pars compacta show Nfe2l2 immunoreactivity within Th-positive neurons. **(B)** Semi-quantitative analysis of Nfe2l2 nuclear intensity within nigral dopaminergic neurons. Data are expressed as fold changes (FC) relative to the control value. Error bars are ± SEM. Data were analysed with one-way ANOVA followed by Tukey’s post hoc test. ^#^P<0.001 *vs.* each of the other three groups.

### GCase inhibition hinder the microglial-mediated induction of NFE2L2 signal in neurons

To characterize the mechanism of GCase inhibition, we investigated the effect of CBE in neuronal cells and in microglia/neuron co-cultures. Initial experiments were carried out in SK-N-BE neuroblastoma cells (Biedler *et al.*, 1978) stably transfected with the bioluminescent reporter of NFE2L2 activity pARE-*luc2*-ires-tdTomato (Rizzi *et al.*, 2017), named SK-ARE-*luc2*. Cells were treated with 200 μM CBE or vehicle for 48 hours in order to ensure the inhibition of GCase activity (Supplementary Fig. 1B), with negligible effects on the activity of additional glycosidase targets (Kuo *et al.*, 2019); 24 hours before harvest, cells were also treated with tBHQ at different concentrations (5 and 15 μM). The enzymatic quantification of luciferase in protein extracts demonstrated a concentration-dependent increase of reporter activity in tBHQ treated cells; however, in contrast with the *in vivo* observation (Fig. 1), the CBE treatment did not affect the NFE2L2 activation induced by tBHQ (Fig. 3A), suggesting that GCase impairment in neuronal cells does not modulate the NFE2L2 activity. Previous data showed that brain cells other than neurons concur in the regulation of oxidative stress responses in the Parkinson’s disease-associated neurodegeneration (Rojo *et al.*, 2010; Rizzi *et al.*, 2018) and may thus provide clue signals that in turn induce the NFE2L2 pathway in neuronal cells. To this aim, we used microglial BV-2 and glioma C6 cell lines to perform co-culturing experiments with SK-ARE-*luc2*. Interestingly, the presence of BV-2 cells per se, but not C6 cells, increased neuronal NFE2L2 activity, an effect also induced by other cells of the macrophage/monocyte lineages (RAW 264.7, mouse and human primary macrophages), but not other lineages (MCF-7) (Fig. 3B). Then we experimentally determined the optimal ratio between BV-2 and SK-ARE-*luc2* to produce the highest NFE2L2 activation; the optimal composition of 1:20 provided the highest level of NFE2L2 activity (+60%), a ratio that more closely represents the physiological neuron/microglia ratio in the brain (Lawson *et al.*, 1990; Bartheld *et al.*, 2016), (Supplementary Fig. 3). The effect was not due to peculiar features of transformed cell lines since an increased NFE2L2 activity of 250% was also observed in co-cultures of primary mouse microglia and neurons, the latter obtained from the ARE-*luc2* reporter mouse, thus carrying the luciferase reporter system for the NFE2L2 pathway (Fig. 3H). These data show that microglia/macrophages communicate with neuronal cells to induce neuronal NFE2L2 activity. Next, we tested the effects of specific inhibition of microglia GCase on neuronal NFE2L2 activity: indeed, treatment with 200 μM CBE, while did not significantly reduce the basal level of NFE2L2 activity (Fig. 3A, 3G), significantly inhibited the microglia-mediated induction of the neuronal NFE2L2 pathway in both transformed lines (−10%) (Fig. 3D) and primary cells (−20%) (Fig. 3I). The observed inhibitory effect was greater in primary cultures than in cell lines, indicating that the microglia-induced activation of the neuronal NFE2L2 pathway is more effective in naïve cells. Finally, these results were confirmed by the semi-quantitative analysis of the fluorescence intensity in immunohistochemistry experiments with an anti-NFE2L2 antibody showing an increased nuclear localization of NFE2L2 in SK-N-BE cells when co-cultured with the microglial cell line, an effect less marked when co-cultures were treated with CBE (Fig. 3E, Supplementary Fig. 4).

**Figure 3.**
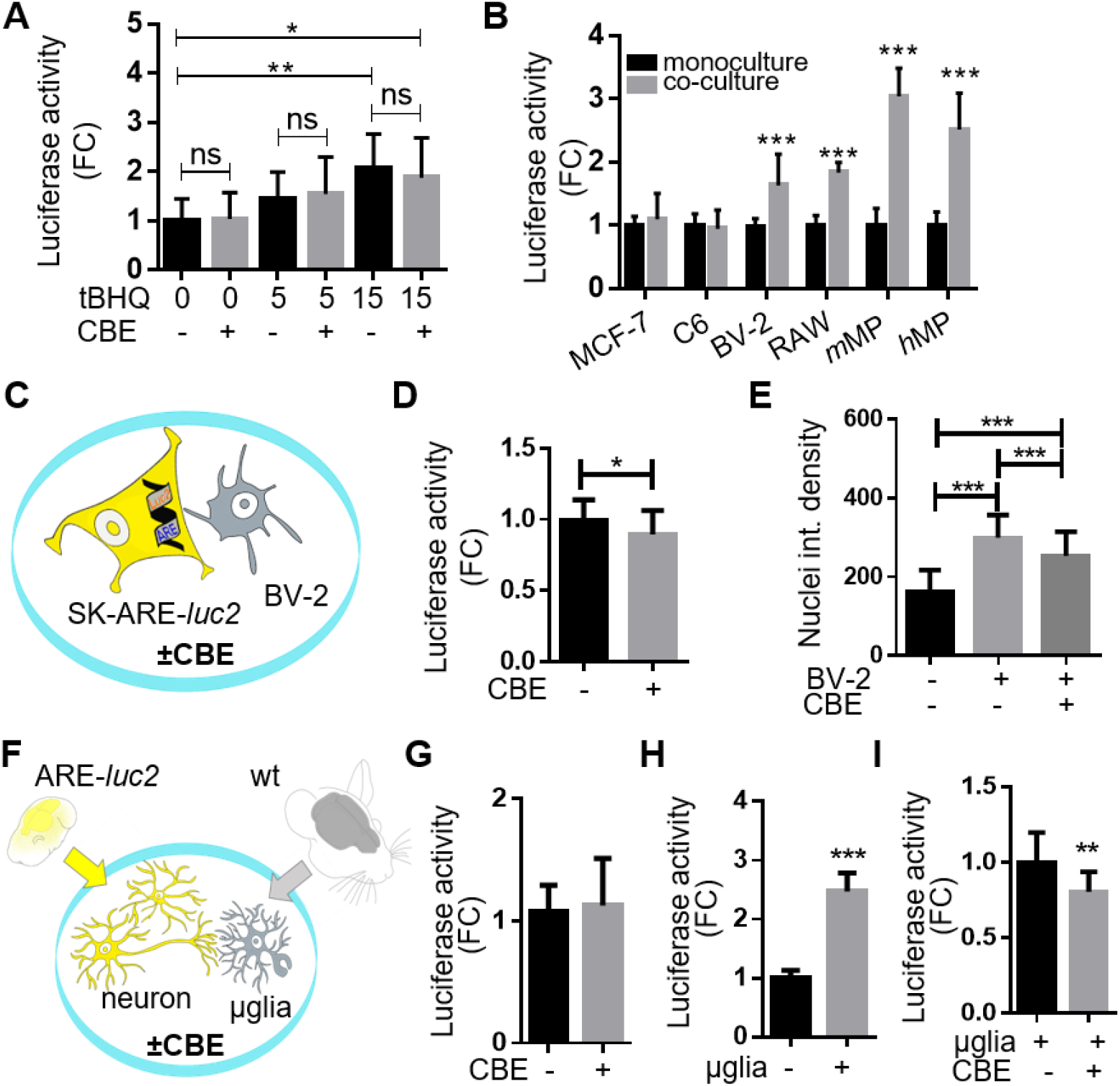
CBE treatment reduces the neuronal NFE2L2 response induced by microglia. (**A**) The NFE2L2 activity was measured in SK-ARE-*luc2* cells treated with 200 μM CBE or vehicle (water) for 48 hours and with 5, 15 μM of tBHQ or vehicle (water) for 24 hours. Luciferase activity is expressed as Relative Light Unit (RLU) for μg of protein extracts normalised *versus* vehicle, bars are mean values ± SD of n=4 measures in triplicate. \**P*<0.05, \*\**P*<0.01 calculated by one-way ANOVA followed by Dunnet’s multiple comparison test *versus* vehicle; ns: not significant by unpaired *t*-test *versus* vehicle. (**B**) The NFE2L2 activity was measured in SK-ARE-*luc2* cultured for 48 hours with MCF-7, C6, BV-2, RAW 249.7 (RAW), murine peritoneal macrophages (mMP), human macrophages (hMP). RLU reported in the graph are FC *versus* monoculture of SK-ARE-*luc2* cells; bars are mean values ± SD for n=3-9 measures in triplicate. ***p<0.001 by 2way ANOVA followed by Sidak’s multiple comparisons test. (**C**) Schematic representation of the experiments reported in **(D-E)** to assess the effect of CBE on neuronal NFE2L2 activation in SK-ARE-*luc2* and BV-2 co-cultures. (**D**) Luciferase activity measured after 48 hours treatment with 200μM CBE or vehicle; RLU values were normalized *versus* vehicle. Bars are mean values ±SD of n=9 measures in triplicate. \**P*<0.05 \*\*\**P*<0.001 calculated by unpaired *t*-test. (**E**) Data derived from the semi-quantitative immunofluorescence analysis of nuclear NFE2L2 staining (Supplementary Fig. 3) on co-cultures treated with 200μM CBE or vehicle for 48 hours. Data are mean values ± SD of the analysis of N=6000 nuclei measured in 20 fields for each condition. \*\*\**P*<0.001 calculated by one-way ANOVA followed by Tukey’s multiple comparison test. (**F**) Schematic representation of the experiments reported in (**H-I**) testing the effects of CBE in primary co-cultures of microglial (from C57Bl/6) and neuronal cell (from ARE-*luc2* reporter mice). (**G**) Luciferase activity from ARE-*luc2* primary neurons treated with 200μM CBE for 48 hours in monoculture, co-cultivated with wild type primary microglia treated with vehicle (**H**) or with 200μM CBE for 48 hours (**I**). RLU values are expressed as FC *versus* vehicle treated samples (**G**), *versus* neuron monoculture (**H**) and *versus* vehicle treated co-cultures (**I**). Data are mean values ±SD of n=5 measures in triplicate. \**P*<0.05, \*\*\**P*<0.001 calculated with the unpaired *t*-test.

### Microglial GCase is essential for the efficient activation of the NFE2L2 pathway in neurons

To corroborate the hypothesis that the CBE-mediated downmodulation of the neuronal NFE2L2 activity observed *in vivo* could be ascribed to the microglial GCase inhibition, either BV-2 or SK-ARE-*luc2* cells were pre-treated with 200 μM CBE for 48 hours before seeding the co-culture (Fig. 4A). Luciferase activity in protein extracts decreased significantly (−25%) only when BV-2 were pre-treated with CBE (Fig. 4B, 4C), demonstrating that inhibition of microglial GCase was necessary and sufficient to reduce the neuronal NFE2L2 activity in the co-culture. This conclusion was confirmed in primary cells i.e. microglia obtained from mice previously i.p. injected with 100 mg/kg/die CBE for 3 days or vehicle (PBS) cultivated for 24 hours on primary neuronal cells derived from ARE-*luc2* mice (Fig. 4D, 4F): microglia obtained from CBE treated mice showed halved GCase activity (Fig. 4E) and a reduction of about 20% of the NFE2L2 activation in primary ARE-*luc2* neurons (Fig. 4F), compared to microglia co-cultures obtained from vehicle-treated mice. The key role played by microglia in the neuronal NFE2L2 activation was firmly demonstrated through pharmacological depletion experiments using PLX3397, a Colony Stimulating Factor-Receptor 1 (CSF-1R) inhibitor (Elmore *et al.*, 2014). Depletion of microglia (Villa et al., 2018) in the ARE-*luc2* mice was obtained after seven intranasal daily administrations of 100 μg PLX3397, twice a day, and half of the treated mice received also two i.p. doses of 75mg/kg tBHQ, 20 and 16 hours before the end of the experiment to induce the NFE2L2 pathway (Fig. 5A). *In vivo* and *ex vivo* imaging quantification of the photon emission from the head area and from the dissected brains of the ARE-*luc2* mice (Fig. 5B, 5C) showed that depletion of microglia hindered the induction of the Nfe2l2 pathway observed upon tBHQ administration (Fig. 5D, 5E). Taken together, these data demonstrated that the neuronal Nfe2l2 response in the brain requires the presence of a functional microglia.

**Figure 4.**
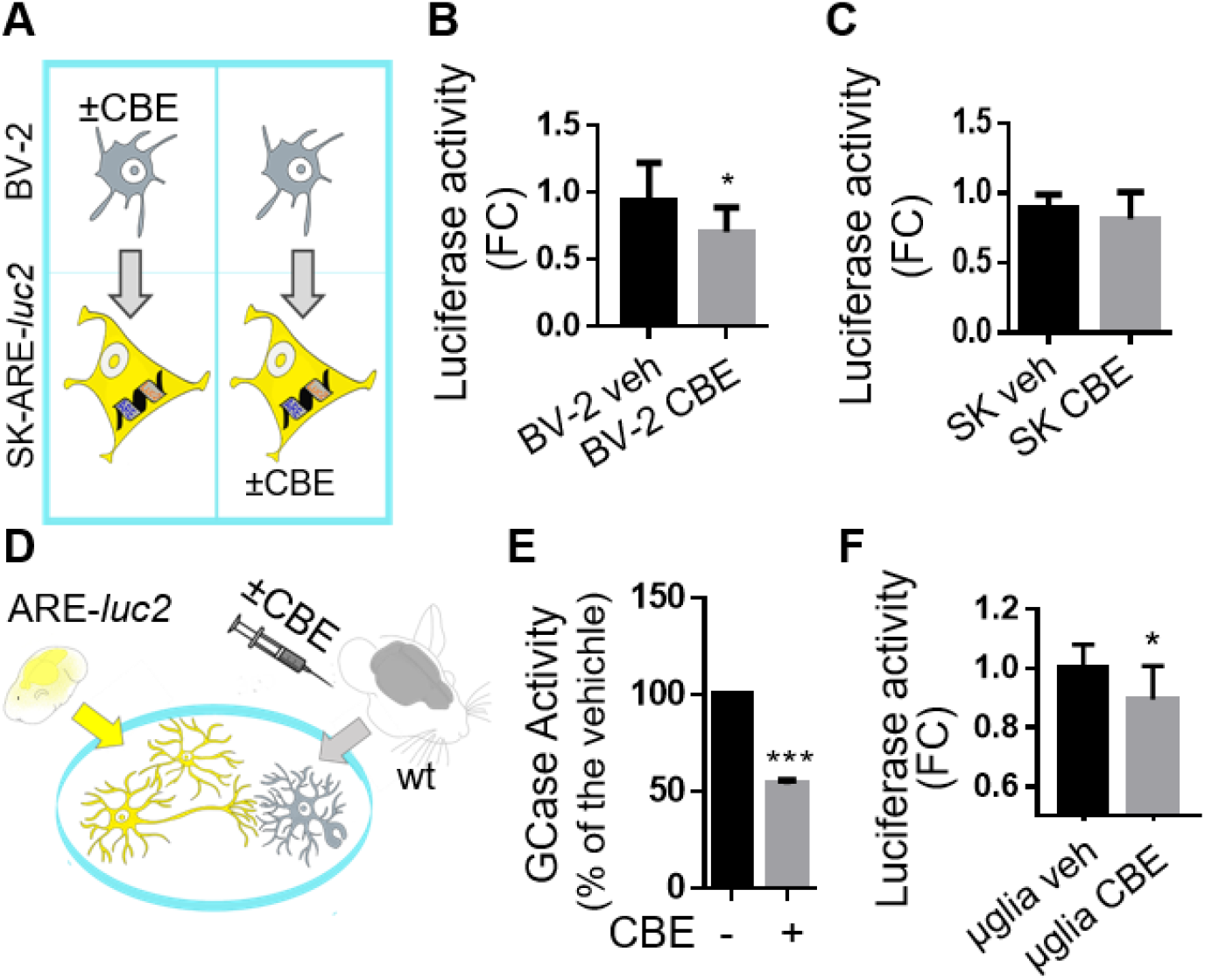
Inhibition of microglial GCase hamper the neuronal Nfe2l2 response. (**A**) Schematic representation of the experiments summarized in (**B, C**) to assess the effect of the selective GCase inhibition in microglial or neuronal cells. BV-2 (**B**) or SK-ARE-*luc2* (**C**) were pre-treated with 200 μM CBE for 48 hours before seeding the co-cultures; graphs report the luciferase activity as FC on vehicle. Data are mean values ± SD of n= 4 measures in triplicate. \**P*<0.05 by unpaired *t*-test. (**D**) Schematic representation of the experiments reported in (**E, F**) aimed at testing the effect of primary microglia extracted from mice treated with CBE 100mg/kg/die for 3 days. (**E**) GCase activity in the primary microglia expressed as % of the activity detected in microglia extracted from the vehicle treated animals. Data are mean values ± SD of n=2 in duplicate. \*\*\**P*<0.001 calculated by unpaired *t*-test. (**F**) Luciferase activity measured in protein extracts derived from ARE*-luc2* neurons co-cultured with microglia derived from CBE or vehicle treated mice. Data are mean values ±SD of n=4 measures in duplicate. \**P*<0.05 calculated by unpaired *t*-test.

**Figure 5.**
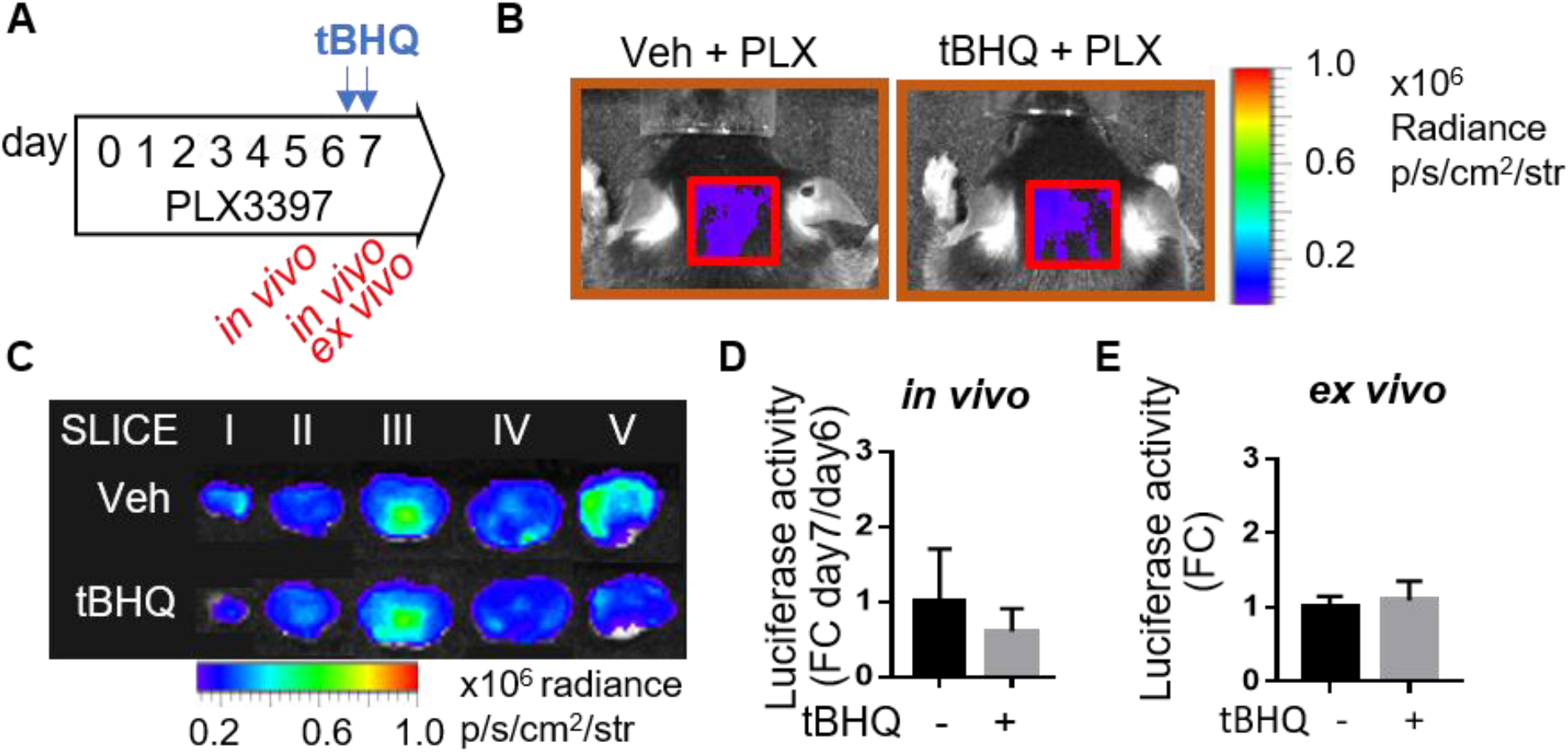
Nrf2 response *in vivo* is microglia-dependent. (**A**) Schematic representation of the pharmacological depletion of microglia by treating mice with 100 μg PLX3397 every 12 hours for 1 week: 20 and 16 hours before the end of the experiment mice received two i.p. injection of tBHQ 75 mg/kg. BLI from the brain area (ROI Fig. 1B) or from the dissected brain acquired for each mouse by *in vivo* (**B**) and *ex vivo* imaging (**C**) are expressed as FC on the value obtained before tBHQ administration (day 6) (for the *in vivo* imaging) (**D**) or on the value of the vehicle treated mice (for *ex vivo* imaging) (**E**). Data are mean values ± SD of n=3 measures. Differences were not significant, by unpaired *t*-test.

### GCase inhibition hampers neuroprotective functions of microglia

The NFE2L2 pathway is of key importance in the detoxification from oxidative insults, thus, we tested whether the microglia-induced activation of the neuronal NFE2L2 pathway was sufficient to increase the detoxification capability of neurons. The microglia-induced neuroprotection was tested in SK-N-BE neuroblastoma cells treated with 0.5 mM MPP+, the active metabolite of MPTP that blocks the mitochondrial electron transport enzyme NADH:ubiquinone reductase (H(+)-translocating) (Nicklas *et al.*, 1985; Vyas *et al.*, 1986) and induces cellular death as a consequence of the formation of reactive oxygen species (Zawada *et al.*, 2011). This treatment increased the cell death of about 50% in the monoculture population of neuroblastoma (Fig. 6A), as quantified by flow cytometry analysis of the dead cells stained with propidium iodide. The cell death was attenuated of about 20% when the neuroblastoma line was co-cultivated with BV-2 microglia cells (Fig. 6B), confirming that microglia was able to increase neuronal resistance to oxidative insults. Conversely, the microglial-dependent neuroprotection was blunted when GCase was inhibited with the 200 μM CBE for 48 hours (Fig. 6D), while the CBE treatment alone did not produce any effect on cell viability in the absence of the neurotoxic stimulation (Fig. 6C). These data demonstrated that GCase inhibition hampers the neuroprotective functions of microglia cells.

**Figure 6.**
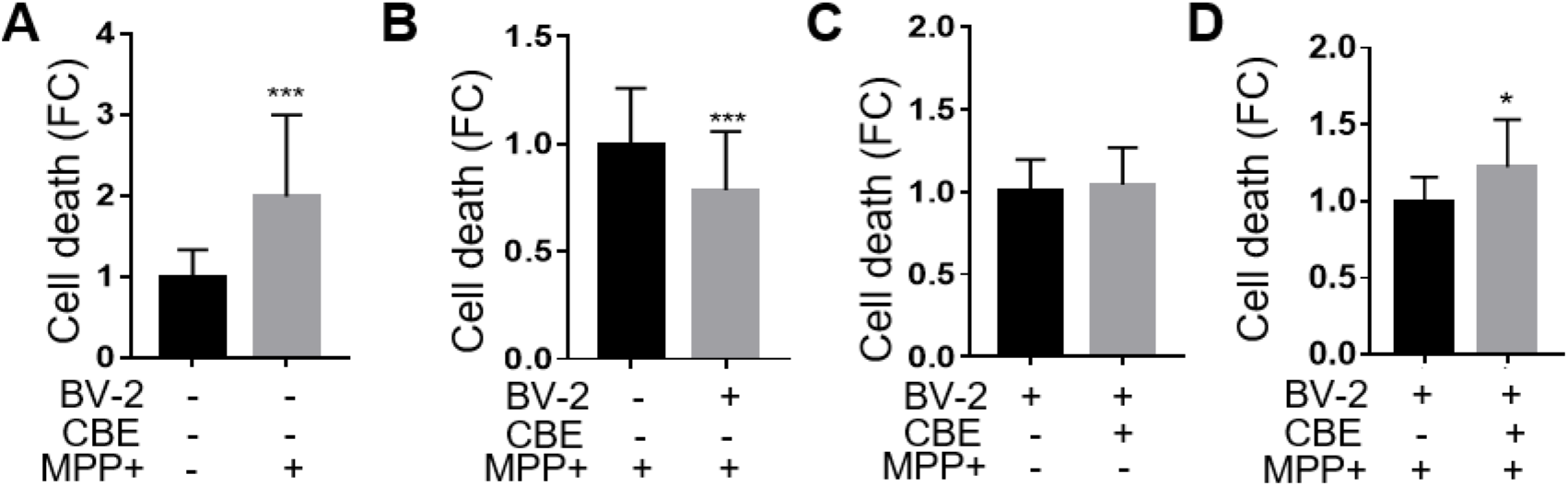
GCase inhibition hampers neuroprotective functions of microglia. Flow cytometry analyses of propidium iodide fluorescence in: (**A**) SK-N-BE treated with 0.5 mM MPP+ or vehicle for 24 hours; (**B**) SK-N-BE cultured alone or in co-culture with BV-2 and treated with 0.5 mM MPP+; (**C**) co-cultures treated with 200 μM CBE for 48 hours or (**D**) with 200 μM CBE for 48 hours and 0.5 mM MPP+ for 24 hours. Data are mean values ±SD of n=6 measures in triplicate of FC *versus* vehicle treated (A, C, D) or monoculture sample (B). \**P*<0.05, \*\*\**P*<0.001 calculated by unpaired *t*-test.

### Inhibition of microglial GCase increases the expression of genes associated with neurodegeneration

To gain molecular insights onto the mechanism of neuroprotection, we measured the expression of genes previously associated with specific microglial phenotypes by real-time PCR analysis on mRNAs purified from BV-2 cells treated with 200 μM CBE or vehicle for 48 hours (Fig. 7). The analysis did not show any increased expression of the proinflammatory cytokine Interleukin 1 beta (*Il-1b*), neither we observed altered regulation of Nfe2l2 target genes (*Nqo1* and *Nfe2l2*) confirming that the GCase inhibitions did no trigger an inflammatory phenotype (Allan *et al.*, 2005; Henkel *et al.*, 2009), neither induce variations in the regulation of the antioxidant and detoxification response in microglia. Surprisingly, the expression of genes involved in the lysosomal pathway (*Lamp1* and *Tfeb*) as well as the expressions of *Cx3cr1*, *C3* and *P2ry12* involved in intercellular communication and selective phagocytosis (Kettenmann *et al.*, 2011; Butovsky *et al.*, 2014), were unaffected suggesting that microglia maintained their homeostatic functionality (Moore *et al.*, 2014; Franco-Bocanegra *et al.*, 2019). Interestingly, two different genes involved in the anti-inflammatory and repairing pathways, *Arg1* and *Trem2* (Benitez and Cruchaga, 2013; Zhao *et al.*, 2018), were upregulated following microglial GCase inhibition. In particular, the latter, involved also in microglia metabolic reprogramming, represents a disease associated gene which mutations are linked with the onset of neurodegenerative diseases such as Alzheimer’s (Karch *et al.*, 2014) and Parkinson’s diseases (Rayaprolu *et al.*, 2013). Recent reports proposed Trem2 as the principal regulator triggering a specific microglial phenotype associated with neural diseases by regulating microglia ability to phagocyte debris (Keren-Shaul *et al.*, 2017).

**Figure 7.**
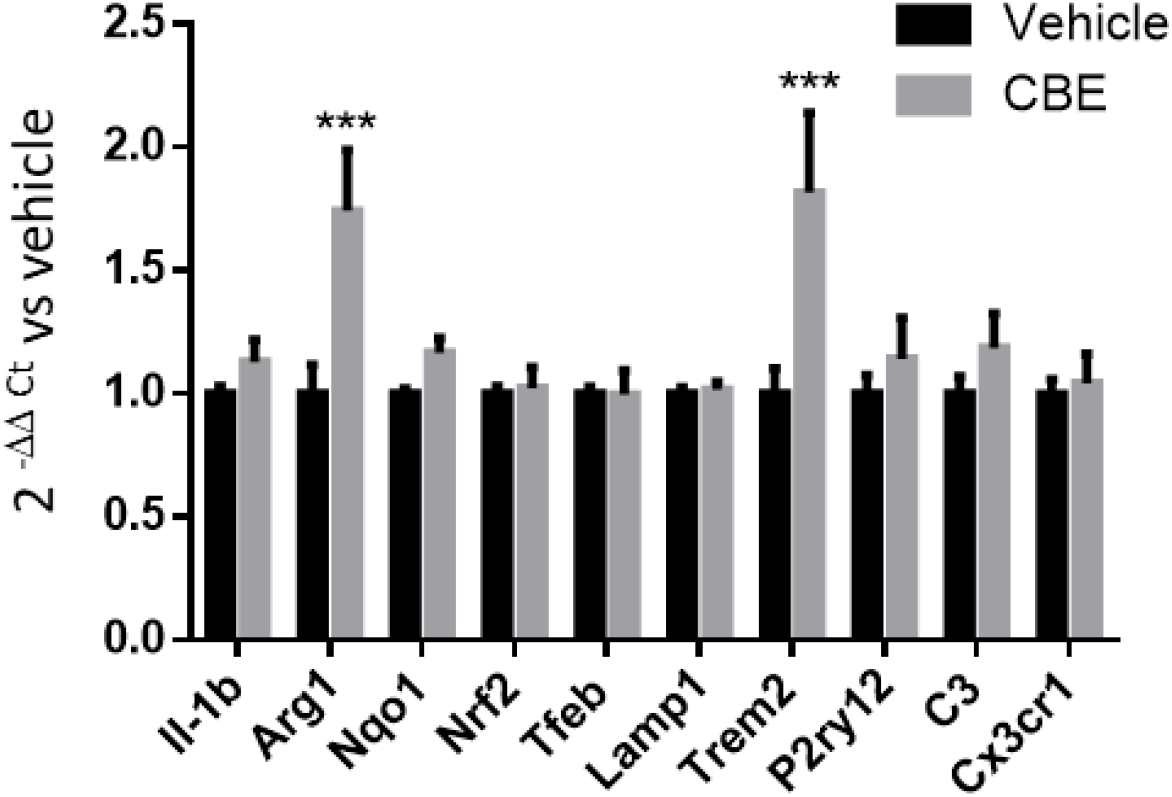
Modulation of gene expression by GCase inhibition in BV-2 cells. Total RNA was purified from BV-2 cells treated with 200 μM CBE for 48 hours and the expression of selected mRNA were analysed by semiquantitative real time PCR. Transcripts were quantified 2^-ΔΔCt^ method. Data are mean values ± SEM for n=3 measures in triplicate. \*\*\**P*<0.001 by two-way ANOVA followed by Sidak’s multiple comparisons test.

These data suggest that GCase may represent a key node in the regulation of the microglia neuroprotective activity ensuring the activation of detoxification/oxidative stress pathways in neurons and modulating a reparative/anti-inflammatory microglial response counteracting neurodegenerative insult.

### Microglia require functional actin-dependent structures to induce NFE2L2 in neuronal dopaminergic cells

Finally, we investigated whether the microglia-to-neuron communication inducing NFE2L2 activity required a paracrine mechanism based on the release of secreted factors or a physical interaction between cells (Biber *et al.*, 2007; Kerschensteiner *et al.*, 2009; Cserép *et al.*, 2020). To this purpose, BV-2 and SK-ARE-*luc2* cells were grown in separate compartments (using a 0.4 μm membrane) of the same well (Fig. 8A) preventing the physical interactions between the two cell types, while preserving the exchange of soluble factors. In this condition, the neuronal NFE2L2 induction could not be observed (Fig. 8B), suggesting that this response does not require diffusible factors. In order to exclude that the membrane pore of 400 nm could obstruct the diffusion of larger macromolecular complexes (e.g. extracellular vesicles), conditioned media derived from co-culture of BV-2 and SK-N-BE cells were added to SK-ARE-*luc2* cells (Fig. 8C); again, the induction of neuronal NFE2L2 activity could not be observed (Fig. 8D), thus definitely ruling out a possible role of microglia secreted factors in this phenomenon. These experiments prompted us to test the alternative hypothesis of a mechanism based on the direct contact between the two cell lineages. Microglia use two major types of branches, strictly regulated by distinct intracellular signaling cascades, to sensing the environment: the actin-dependent filopodia and the tubulin dependent large processes (Nolte *et al.*, 1996; Nimmerjahn *et al.*, 2005; Bernier *et al.*, 2019; Cserép *et al.*, 2020). To define which of the two pathways is required for the microglia-dependent induction of the NFE2L2 activity, BV-2 cells were pretreated with nocodazole, an agent preventing microtubule assembly, at a concentration and time span (25 μM for 2 hours) known to inhibit large process without affecting filopodia movement (Bernier *et al.*, 2019). After the treatment, cells were seeded on the SK-ARE-*luc2* cells and luciferase activity was measured 24 hours later: the results demonstrated that pharmacological disruption of microglial large processes did not affect the ability of microglia to induce NFE2L2 signaling in neurons (Fig. 8E). On the contrary, when BV-2 cells where pretreated with 1 μM cytochalasin for 1 hour, a treatment blocking actin polymerization and reducing filopodia motility (Pan *et al.*, 2011; Bernier *et al.*, 2019), we observed a significant decrease (>25%) in the induction of neuronal NFE2L2 activity (Fig. 8F) suggesting that actin-dependent structures are necessary for the microglia-mediated stimulation of NFE2L2 pathway in neurons.

**Figure 8.**
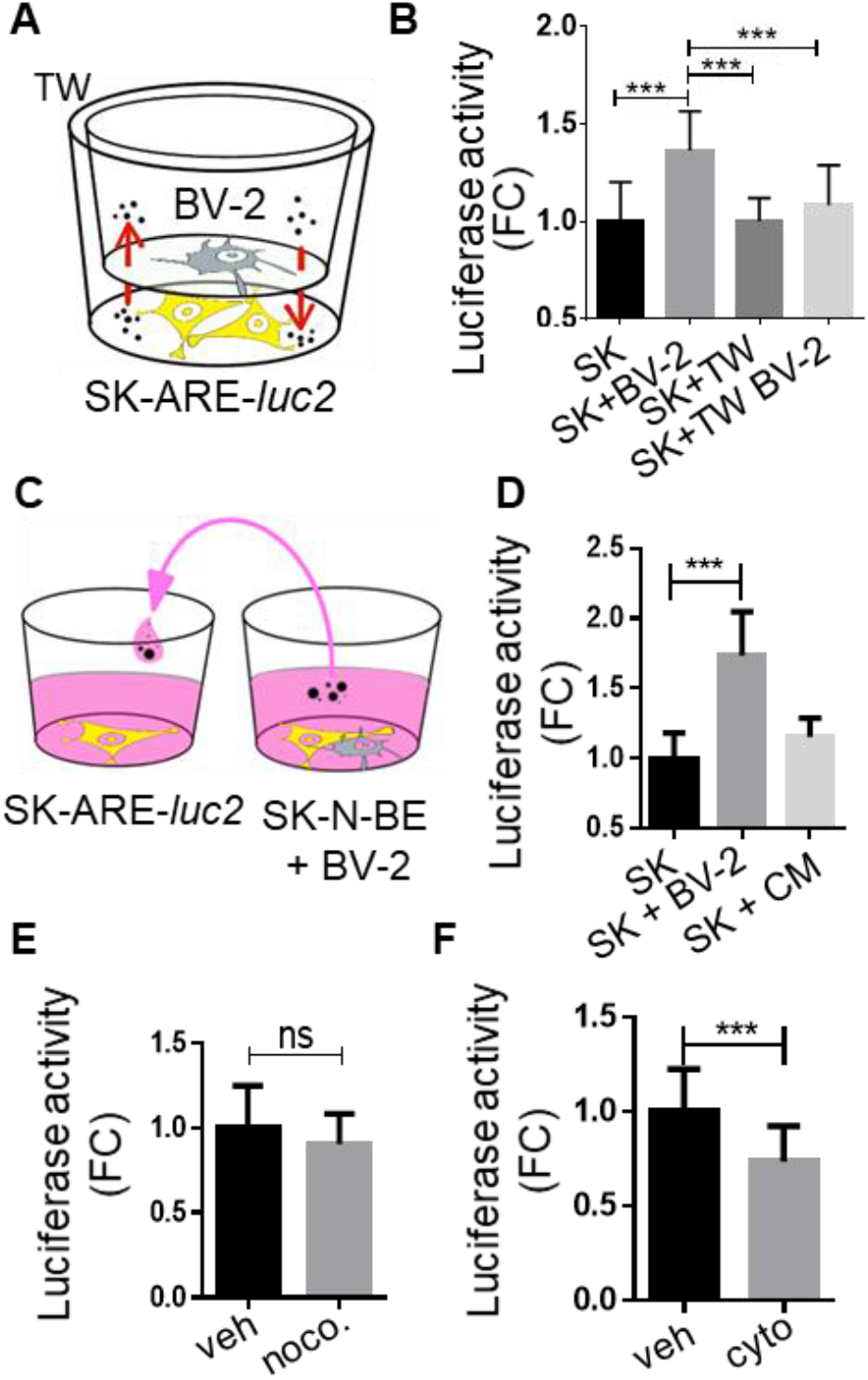
Mechanism of microglia-to-neuron cross-talk. (**A**) Scheme of the transwell (TW) tool used in the experiment reported in (**B**): BV-2 and SK-ARE-*luc2* cells were seeded in separate compartment of the TW. B) Luciferase enzymatic activity obtained from co-culture of SK-ARE-*luc2* and BV-2 cells (SK+BV-2), or SK-ARE-*luc2* growth alone in the transwell (SK+TW), or growth in separated compartment with microglia (SK+TW+BV-2); values are expressed as RLU normalized *versus* monoculture of SK-ARE-*luc2* (SK). Data are mean values ±SD of n=6 measures in triplicate. \*\*\**P*<0.001, by one-way ANOVA followed by Tukey’s multiple comparison test. (**C**) Schematic representation of the experiment reported in (**D**): SK-ARE-*luc2* cells were grown in co-culture with BV-2 (SK+BV-2) or in monoculture with the conditioned medium (SK+CM) obtained from the co-culture. (**D**) Luciferase activity (RLU) is expressed as FC *versus* SK-ARE-*luc2* monoculture (SK). Data are mean values ±SD of n=3 measures in triplicate. \*\*\**P*<0.001, by one-way ANOVA followed by Dunnet’s multiple comparison test *versus* SK. (**E**) BV-2 were pre-treated with 25μM nocodazole (noco) or vehicle for 2 hours and co-cultured SK; Luciferase activity (RLU) are FC *versus* vehicle (veh); data are mean values ±SD of n=12 measures in quadruplicate. Mean differences are not significant as calculated by unpaired *t*-test. (**F**) BV-2 were pre-treated with 1μM cytochalasin (cyto) or vehicle (veh) for 1 hour and co-cultured with SK-ARE-*luc2*. Luciferase activity (RLU) are FC *versus* vehicle: data are represented as mean ±SD for n=8 measures in triplicate.

## Discussion

The neurodegenerative phenotype promoted by GBA mutations has been so far attributed to the loss of function of the GCase enzyme in neurons, although microglia could be another plausible target cell type, since the macrophage lineage is responsible for the peripheral effects of these mutations (Prinz *et al.*, 2014). So far, most of the investigation have focussed on neurons mainly because it is currently believed that the progressive accumulation of the GCase substrate may interfere with macrophage lysosomal activity only when both GBA alleles are mutated (Grabowski and Beutler, 2001). However, considering that GBA is highly expressed in microglia (Zhang *et al.*, 2014) and mutations are able to significantly reduce the GCase enzymatic activity (Beutler, 1992), we hypothesized that even a partial inhibition of this enzyme could be sufficient to produce functional consequences in microglia physiology. Indeed, we here demonstrate that the microglia-dependent activation of the neuronal NFE2L2 activity and the functional consequences of GCase inhibition in microglia affect the neuronal redox response and cell survival to neurotoxicity. This mechanism requires a direct contact between the two cell lineage and functional actin-dependent structures, namely the cytoskeleton and filopodia, which are required for the motility of microglial cells (Bernier *et al.*, 2019; Cserép *et al.*, 2020). We may assume that the cell-to-cell contact could be hindered by the GCase inhibition, as suggested by recent observations from our lab showing that CBE treatment induces a round shaped morphology in a subpopulation of microglia cells (Supplementary Fig. 5), that, by reducing cell protrusions, might curtail their ability to interact with neurons and induce the NFE2L2 pathway. The consequence of the reduced NFE2L2 activity is an increased susceptibility of neuronal cells to oxidative stress or toxic molecules; in keeping with this conclusion, our co-culture experiments demonstrate that reduced GCase activity is associated with a higher sensitivity of dopaminergic neurons to MPP+ treatments (Fig. 6). Interestingly, the microglial phenotype associated with GCase inhibition seems to be more related with the anti-inflammatory phenotype, since we showed upregulation of anti-inflammatory genes involved in reparative responses (*Arg1* and *Trem2*), possibly because of the activation of a cell protection mechanism. We may speculate that in early neurodegenerative stages, anti-inflammatory microglia may prevail over the pro-inflammatory phenotype that has been observed late in the neurodegenerative process, likely as a consequence of the extensive progression of neuronal death (Vitner *et al.*, 2012). Future studies will clarify the relation of the anti-inflammatory programme stimulated by CBE and the round shaped, poorly communicative phenotype of microglia highlighted in our experiments and its role in neurodegeneration.

Microglia establish a paracrine network of connections with the other brain cells mediated by soluble factors (Kerschensteiner *et al.*, 2009), extracellular vesicles (Paolicelli *et al.*, 2019) or by a direct cell-to-cell contacts (Cserép *et al.*, 2020; Vainchtein and Molofsky, 2020) involving neuronal dendrites or soma; through these connections, specific microglia functions participate in brain homeostasis and neuronal defense in case of brain injury or infections. Our study revealed a novel type of neuroprotection mechanism, based on the direct cell-to-cell contact, which enrich the microglia armamentarium for neuronal protection. We demonstrated that a 50% reduction in microglial GCase activity, recapitulating the enzymatic activity impairment observed in heterozygotic carriers of severe GBA mutations (Sanchez-Martinez *et al.*, 2016), is sufficient to destabilize the neuronal capacity to reacting against oxidative stress conditions or neurotoxic insults. In our experiments, *in vivo* CBE treatment was appropriately titrated (Vardi *et al.*, 2016) to obtain a comparable reduction; as a result of this treatment, we observed a slight, 20% inhibition of neuronal NFE2L2 activation (Fig. 4): the extent of this impairment was proven sufficient to produce significant effects on neuron survival *in vitro* in the presence of a neurotoxic stimulus like MPP+ (Fig. 6). We believe that this vulnerability is especially relevant for dopaminergic neurons, that display high oxidative stress levels and increased susceptibility to neurotoxic stimuli, as a consequence of the dopamine metabolism (Hermida-Ameijeiras *et al.*, 2004); indeed, high oxidative stress levels were observed *post mortem* in the brain of Parkinson’s patients (Zhou *et al.*, 2008). The increased oxidative stress together with the reduction in glutathione content in dopaminergic neurons was associated with a compensatory hyperactivation of the NFE2L2 pathway in SNpc neurons (Toulorge *et al.*, 2016). In these conditions, the slight but constant reduction in NFE2L2 activity induced by GBA mutations in microglia may progressively hinder the full protective response against neurotoxic stimulations. It is thus conceivable that with a reduced NFE2L2 activity, the oxidative stress, at length, may produce negative consequences on cellular viability particularly in dopaminergic neurons. Accumulation of damage due to the continuous impairment of NFE2L2 signal is fitting with the slight, but constant, neuronal death occurring in the long-lasting process of Parkinson’s pathogenesis, starting years before the onset of clinical manifestations (Postuma and Berg, 2016). It is true that not all GBA carriers develop Parkinson’s disease, we may hypothesize that the greater vulnerability of neurons to neurotoxic stimuli in GBA carriers depends on the individual variability of exposition to environmental factors (neurotoxic stimuli) and/or on different levels of oxidative stress associated with genetic factors: both conditions may account for the different penetrance of specific GBA mutations in the development of the neurodegenerative phenotype.

In conclusion, a large number of studies highlighted the neuronal effects of GBA mutations on lysosomal and mitochondrial neuronal functions, leading to increased alpha-synuclein deposition and vulnerability to neurotoxic insults; in our study, we demonstrated that GCase inhibition impairs also microglial ability to induce a detoxification response through the modulation of the NFE2L2 signalling pathway in neurons. Alterations in this protective mechanism could increase the risk of neurodegeneration induced by toxic molecules and oxidative stress, especially important for dopaminergic neurons characterized by an elevated oxidative metabolism. Unveiling this mechanism might contribute to identify novel targets to restore microglial neuroprotective functions and prevent the onset of neurodegeneration in Gaucher’s and Parkinson’s diseases.

## Methods

### Reagents

All reagents were purchased from Merck. tert-butylhydroquinone (tBHQ, Cat. 112941), conduritol-B-epoxide (CBE, Cat. 234599), cytochalasin D (Cat. C8273), nocodazole (Cat. M1404), PLX3397 (Cat. S7818-50, DBA).

### Cell culture

All cell lines were purchased from the American Type Culture Collection ATCC. Primary neurons where derived from neural tissue following standard operational procedure from p0-1 mice brain using the Neural Tissue Dissociation Kit-Postnatal neurons (Cat. 130-094-802, Miltenyi Biotec). After removing the meninges, brains cortex from six mice were pooled as a single experimental group and subjected to an enzymatic and mechanical dissociation at 37°C, the cellular suspension where filtered with 70 μm strainer and seeded on poly-L-ornithine-coated plates. Half of medium volume was replaced each 2 or 3 days. Microglia were isolated from whole brains of adult (age 3-6 months) male mice with a previously described protocol (Villa *et al.*, 2018). Brains from two mice were pooled and subjected to an enzymatic and mechanical dissociation, after myelin removal, cells were processed for microglia magnetic sorting with magnetic column to purify CD11b+ cells, namely microglia (Cat. 130-093-634, Miltenyi Biotec). For generation of human blood-derived macrophages we followed a previously described procedure (Toniolo *et al.*, 2015), human peripheral blood mononuclear cells were isolated from buffy coats obtained from healthy volunteers (Niguarda Hospital) after Ficoll®Paque Plus (Cat. 17-1440-02, GE Healthcare) density gradient centrifugation. Before the use, peripheral blood mononuclear cells were growth for 1 week in DMEM media (Cat. 32430-027, Gibco) supplemented with 10% FBS (Fetal Bovine Serum Ultra-Low Endotoxin, ECS0186L, Euroclone), 1% streptomycin–penicillin (Cat. 15240-062, Gibco), 1 % GlutaMAX™ (Cat. 35050-061, Gibco), 50 ng/mL of M-CSF (Cat. 11343113, Immunotools), and media were exchanged every 2 days. Primary peritoneal murine macrophages were extracted as described in (Mornata *et al.*, 2020). Three mice of 3-6 months were euthanized and subjected to a peritoneal lavage with PBS to extract macrophages subsequently purified using CD11b+ magnetic beads (Cat. 130-049-601, Miltenyi Biotec), following the operative manual. Human macrophages were detached from the plate using StemPro™ Accutase™ (Cat. A11105, Gibco), centrifuged and resuspended in RPMI complete media. SK-ARE-*luc2* cells were obtained by stable transfection of SK-N-BE cells. Briefly, cells were transfected with pARE-*luc2*-ires-tdTomato (Rizzi *et al.*, 2018) and pSTC1-Neo DNA plasmids (10:1 ratio) using Lipofectamine LTX & PLUS reagent (Cat. 15338, Thermo Fisher Scientific), with a DNA:Lipofectamine LTX:PLUS reagent ratio of 2.5:6.5:1.5, and following manufacturer’s instructions. Forty-eight hours after transfection, cells were re-seeded at different concentrations and positive clones identified by selection using 300 μg/mL G418 (Cat. G8168, Sigma Aldrich) in RPMI 1640 (Cat. 61870044, Gibco) plus 10% fetal calf serum and 1 mM Sodium pyruvate (Cat. 11360, Thermo Fisher). Clones were assayed for their ability to respond to tBHQ, a well-known NFE2L2 activator, by increasing luciferase production (Supplementary Fig. 1). For co-cultures experiments, 150,000 primary neuronal cells or 70,000 SK-N-BE or SK-ARE-*luc2* cells were plated in each well of a 24-well plate and cultured for 10 days (primary cells) or 1 day (cell lines). Then, primary microglia, BV-2, MCF-7, RAW 264.7 or primary macrophages were seeded over the neuron layer: if not otherwise specified, the number of seeded cells were 37,500 cells/well for primary microglia and primary macrophages, or 3,500 cells/well for BV-2, RAW 264.7 and MCF-7. For transwell experiments, 0.4 μm pore polyester membrane inserts (Cat. 3460, Corning) were used to separate BV-2 cells from adhered neuroblast cells. Neuronal and neuronal-microglial co-cultures were grown in Neurobasal A medium (Cat. 10888-022, LifeTechnologies) containing 1% streptomycin–penicillin, 1% GlutaMAX™, 2% B-27™ Supplement (Cat. 17504-044; Gibco), 10 mM HEPES (Cat. H0887, Merk), in a humidified 5% CO2-95% air atmosphere at 37 °C. Other co-cultures were grown in RPMI 1640 Media containing fetal bovine serum to a final concentration of 10%, 1% streptomycin–penicillin and 1% GlutaMAX™.

### Cell treatments

If not otherwise specified, cells were treated with 200 μM CBE or vehicle (water) for 48 hours, with 1μM cytochalasin D or vehicle (0.0005% v/v EtOH final) for 1 hour, with 2 μM nocodazole or vehicle (0.02% v/v EtOH, 0.01% v/v DMSO final) for 2 hours, with 5 and 15 μM tBHQ or vehicle (water) for 24 hours.

### Flow cytometry assay

Flow cytometry experiments were performed on at least 200,000 cells for each sample by using Novocyte 3000 (Agilent Technologies, Inc) equipped with 488 nm lasers. Cells were incubated 2 minutes at 4°C with 0.1 mg/ml propidium iodide (Cat. P4170, Sigma Aldrich) and fluorescence pulses were detected using 585/40 nm collection filter. Results were analysed using NovoExpress software (Agilent Technologies).

### Luciferase enzymatic assay

Luciferase assays was performed as described (Rizzi *et al.*, 2018). Briefly, cells were lysed with Luciferase Cell Culture Lysis Reagent (Cat. E1531, Promega) and protein concentration was determined with Bradford assay (Bradford, 1976); the biochemical assay of luciferase activity was carried out in luciferase assay buffer by measuring luminescence emission with a luminometer (Veritas, Turner Biosystems,) and the relative luminescence units (RLU) determined during 10 seconds measurements.

### Glucocerebrosidase assay

Glucocerebrosidase assay was performed as previously described (Mus *et al.*, 2019). Cells were lysed with RIPA buffer and protein concentration determined with BCA (Cat. 23227, Pierce). The biochemical assay of GCase activity was carried out in a buffer containing 4-methylumbelliferyl beta-d-glucopyranoside substrate (Cat. M3633, Sigma Aldrich); after 1 hour at 37°C the reaction was stopped with 0.25 M glycine buffer pH 10.4 and the fluorescence emission of the 4-methylumbelliferyl generated (4-MU) by the reaction was read with a fluorimeter (EnSpire Plate Reader, PerkinElmer) selecting an excitation wavelength of 365 nm and an emission wavelength of 445 nm. The determined enzyme activity was expressed as μmol 4-methylumbellifery generated in 1 hour per μg of protein and normalized on vehicle-treated cells.

### Immunohistochemical and immunocytochemical analysis

Mice were sacrificed with an i.p. injection of sodium pentobarbital and transcardially perfused with cold saline solution followed by 4% paraformaldehyde in PBS. Brains were immediately removed, post-fixed in 4% paraformaldehyde for 24 hours and then cryopreserved in 30% sucrose. Serial coronal sections of 35 μm were cut throughout the brain using a freezing microtome. Free-floating sections were rinsed in Tris-HCl (pH 7.6), and non-specific binding sites were blocked by incubation in Tris-buffered saline solution containing 0.25% Triton-X-100 and 5% normal horse serum at room temperature for 1 hour. Sections were kept for two nights at 4 °C in Tris-buffered saline with 1% BSA, 0.25% Triton-X-100 and the following primary antibodies: mouse anti-tyrosine hydroxylase (Th) antibody (1:300; Cat. MAB318, Chemicon) and rabbit anti-Nfe2l2 antibody (1:300; Cat. AB3116, Abcam). They were then rinsed and incubated with horse anti-rabbit secondary antibody Dylight 594 (1:300, Vector Laboratories) and horse anti-mouse antibody Dylight 649 (1:300, Vector Laboratories) for 1 hour at 4 °C. Samples were finally counterstained with DAPI (1:10,000 in Tris-HCl, Thermo Fisher Scientific), mounted with a coverslip using Vectashield mounting medium (Vector Laboratories). Fluorescence stack images were collected at 1.29 μm intervals throughout the midbrain using a Zeiss microscopes (LSM880) with a 40× Plan-Apochromat objective. Semi-quantitative analysis of Nfe2l2 nuclear intensity was performed within neurons of the substantia nigra pars compacta (SNpc). For each animal, two SNpc-containing midbrain sections were used for this analysis. Sections were collected at similar anatomical levels and processed with Fiji software (ImageJ, NIH, version 2.0.0). The software selected and delineated DAPI-stained nuclear areas and, within these areas, Nfe2l2 grey values were measured and averaged.

Co-cultures of SK-N-BE and BV-2 cells were fixed in 4% paraformaldehyde fixative for 15 minutes and washed with PBS. The fixed cells were incubated at room temperature for 1 hour in blocking solution containing 0.1% p/v BSA (Cat. A9418, Merk), 10% v/v Goat serum (Cat. ECS0200D, Euroclone), 0.1% v/v Triton-X100 (Sigma, T-9284). Next, cells were incubated at 4°C with 10% blocking solution in PBS with a rat anti-CD11b antibody (diluted 1:500; Cat. MCA7114, Serotec) and a rabbit anti-NFE2L2 antibody (diluted 1:500; Cat. C-20, Santa Cruz biotechnology). After 24 hours, fixed cells were rinsed in PBS and incubated 1 hour at RT 10% blocking solution in PBS with a mixture of Alexa Fluor 488 conjugated goat anti-rat IgG antibody (diluted 1:300, Cat. A-11006, Molecular Probes Inc.) and Alexa Fluor 555 conjugated goat anti-rabbit IgG antibody (diluted 1:300; Cat. A-21429, Molecular Probes Inc.). Finally, fixed cells were rinsed in PBS and covered with 1:1000 DAPI solution in PBS (Invitrogen, Cat. D1306). The fluorescence image acquisition was performed using an Axiovert 200M microscope with a dedicated software (AxioVision Rel 4.9, Zeiss), taking 20 random field for condition. The images were elaborated with ImageJ Fiji software, in brief: the ROI of cellular nuclei automatically generated by the software using the field of DAPI were utilized to quantify the average gray value of the NFE2L2 field. The value related to the nuclei of Cd11b+ positive cells were manually excluded from the analysis.

### Animal treatments

All animal experimentation was carried out in accordance with the ARRIVE and European Guidelines for Animal Care. All animal experiments were approved by the Italian Ministry of Research and University or the ethical committee of the State Agency for Nature, Environment and Consumer Protection in North Rhine Westphalia. The animals were fed ad libitum and housed in individually ventilated plastic cages within a temperature range of 22–25 °C, relative humidity of 50% ± 10% and under an automatic cycle of 12 hours light/dark (lights on at 07:00 AM). ARE-*luc2* mice where generated in our lab (Rizzi *et al.*, 2017, 2018). Pharmacological treatments: mice (15–30 weeks old) were i) i.p. injected with 100 mg/kg/die for 3 days of CBE or vehicle (PBS), ii) i.p. injected with tBHQ 75 mg/kg dissolved in PBS + 1%DMSO + 20%PEG300, iii) 100 μg PLX3397 or vehicle solution (5% DMSO + 45% PEG300 + ddH2O) were given as nose drops (3μl/drop in each nostril corresponding to 6 μl/administration), alternating between the left and right nostrils for two times, at intervals of 2 min; subsequent doses were given each 12 h, for 1 week. For the transnasal administration anesthetized mice were placed in supine position, and a heated pad was inserted under the dorsal neck to induce a hyperextension of the head back (Villa *et al.*, 2019).

### *In vivo* and *ex vivo* imaging

For the semi-quantitative analysis of photon emission, animals were injected s.c. with 80 mg/kg of luciferin 15 min prior the imaging session. Mice were anaesthetized using Isofluorane and kept under anesthesia during the 5 min of each optical imaging session carried out with a CCD-camera (IVIS Lumina II Quantitative Fluorescent and Bioluminescent Imaging, PerkinElmer); photon emission in different brain areas was measured using the Living Image Software v. 4.2 (Perkin Elmer). Mice were killed by cervical dislocation after the last *in vivo* acquisition, brains were rapidly dissected and sectioned by means of a “brain matrix” (adult mouse, coronal and sagittal, 1mm spacing). Sections were immediately subjected to *ex vivo* imaging session of 5 minutes. Photon emission was quantified with the Living Image Software v. 4.2.

### Real-time PCR

Real Time RT-PCR analyses were performed as previously described (Rizzi *et al.*, 2018). The following primers were used:

Sequence:

*Rplp0*: 5′-GGCGACCTGGAAGTCCAACT-3′, 5′-CCATCAGCACCACAGCCTTC-3′;

*Nfe2l2*: 5′-CCCAGCAGGACATGGATTTGA-3′, 5′-AGCTCATAGTCCTTCTGTCGC-3′;

*Arg1*: 5′-CAGAAGAATGGAAGAGTCAG-3′, 5′-CAGATATGCAGGGAGTCACC-3′;

*Trem2*: 5′-GGAACCGTCACCATCACTCT-3′, 5′-CTTGATTCCTGGAGGTGCTGT-3′;

*Il-1b*: 5′-TGCCACCTTTTGACAGTGATG-3′, 5′-GCTGCGAGATTTGAAGCTGG-3′;

*Nqo1* 5′-GGTAGCGGCTCCATGTACTC-3′, 5′-CGCAGGATGCCACTCTGAAT-3′;

*Tfeb*: 5′-CCAGAAGCGAGAGCTCACAGAT-3′, 5′-TGTGATTGTCTTTCTTCTGCCGC-5′;

*Lamp1*: 5′-GCCCTGGAATTGCAGTTTGG-3′, 5′-TGCTGAATGTGGGCACTAGG-3′;

*Cx3cr1*: 5′-CTGCTCAGGACCTCACCATG-3′, 5′-CACCAGACCGAACGTGAAGA-3′;

*P2ry12*: 5′-GAACCAGGACCATGGATGTG-3′, 5′-CCAAGCTGTTCGTGATGAGC-3′;

*C3:* 5′-GGAAGATCCGAGCCTTTTAC-3′, 5′-CCACACCATCCTCAATCACTAC-3′

### Statistical analysis

Statistical analyses, unless otherwise indicated in the figure legend, were done using *t*-test, one-way ANOVA or two-way ANOVA for multiple treatment comparisons (Graph Pad 7 software). A *p*-value lower than 0.05 was considered as statistically significant.

### Data availability

The data that support the findings of this study are available from the senior author upon reasonable request.

## Abbreviations

CBE: conduritol-B-epoxide
GCase: β-glucocerebrosidase
MPP+: 1-methyl-4-phenylpyridinium
MPTP: 1-methyl-4-phenyl-1,2,3,6-tetrahydropyridine
NFE2L2: nuclear factor erythroid 2-related factor 2
FC: fold change
SNpc: Substantia Nigra pars compacta
tBHQ: tert-butylhydroquinone
Th: tyrosine hydroxylase

## Acknowledgments

The authors are indebted to Finlombarda and the TOPsrl research team for generating the reporter mouse ARE-*luc2*.

## Supplementary Figures

**Supplementary Figure 1.**
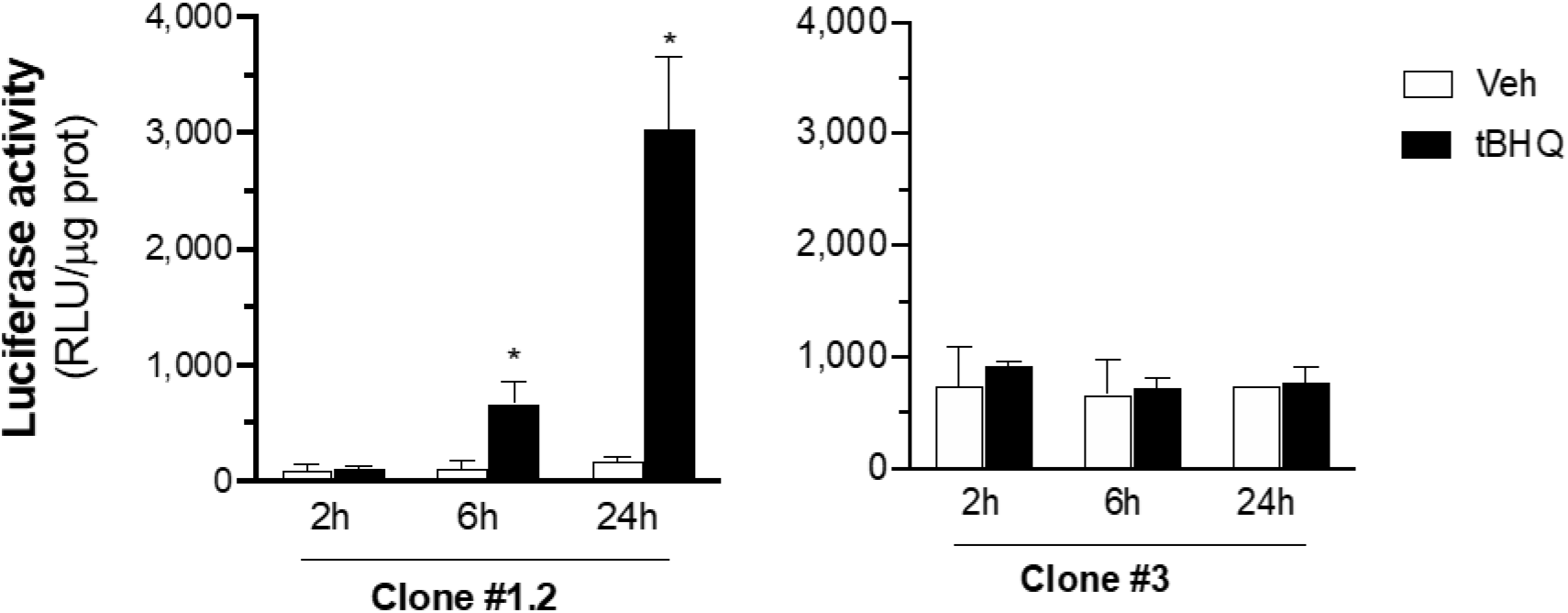
SK-ARE-*luc2* clone selection. Following transfection with the pARE-*luc2*-ires-tdTomato plasmid, SK-N-BE cells were subject for 4 weeks to a selective pressure with G418 and two representative clones #1.2 and #3 tested for the ability to report NRF2 upregulation 2, 6 or 24 hours after the treatment with 80 μM tBHQ or vehicle. Clone #1.2 was selected, amplified and used in the present study. Luciferase enzyme activity expressed as relative luciferase units (RLU) per μg protein, data are mean values ± SEM (n = 3) of a single experiment, which is representative of at least two other independent experiments. \**P*<0.05 vs vehicle calculated by one-way ANOVA followed by Tukey’s multiple comparison test.

**Supplementary Figure 2.**
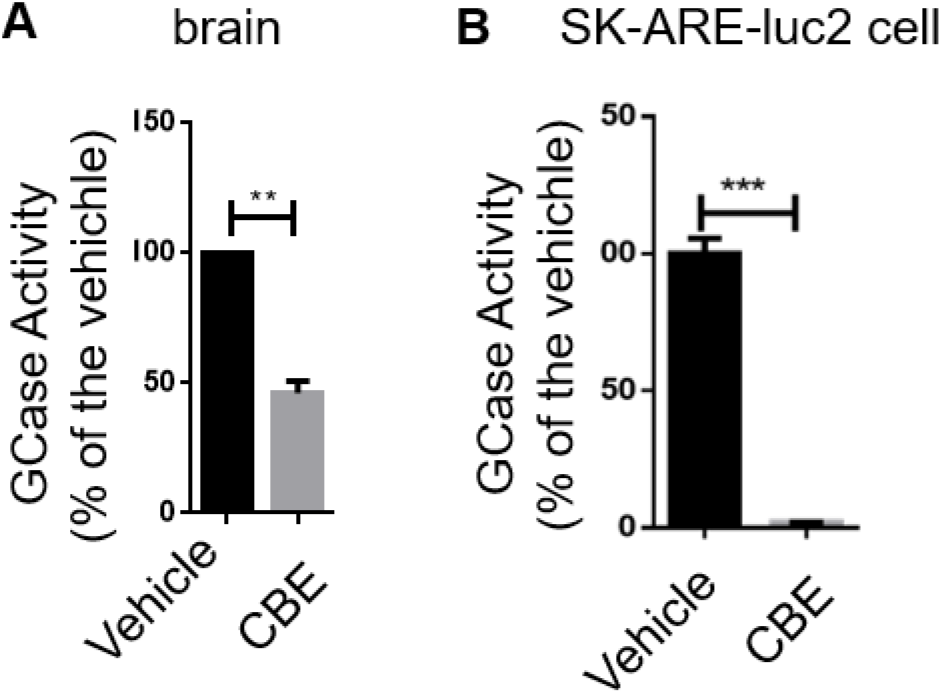
GCase inhibition in mice and SK-ARE-*luc2* cells treated with CBE. (**A**) Residual activity of GCase in the brain of mice treated with 100 mg/kg/ CBE for three days. (**B**) Residual activity of GCase in SK-ARE-*luc2* cells treated with 200 μM CBE for 48 hours. Data are mean values of the enzymatic activity quantified as μmol 4-MU generated in 1-hour reaction per μg of proteins expressed as % of the activity detected versus vehicle treated animals (A) or cells (B) ±SD of n=2 in duplicate (*in vivo* experiments), n=2 in triplicate (cell culture experiments). \*\**P* < 0.01, \*\*\**P*<0.001 *versus* vehicle calculated by unpaired t-test.

**Supplementary Figure 3.**
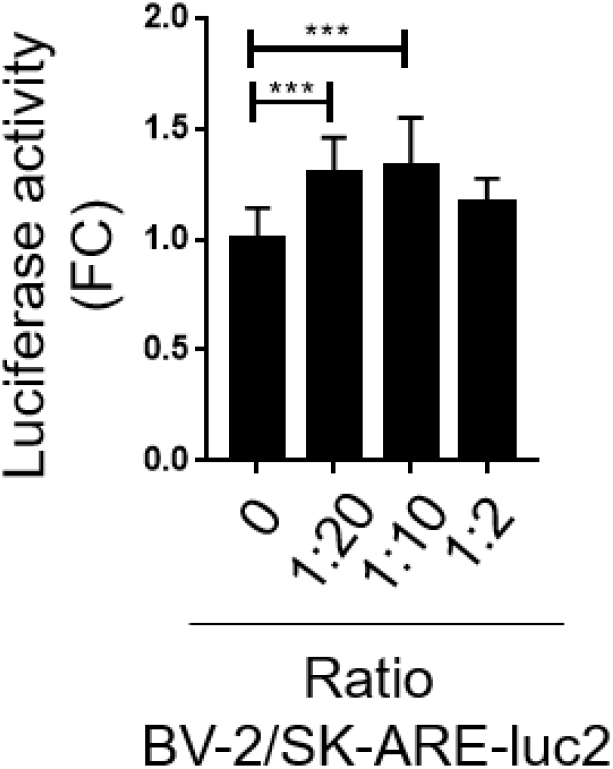
Different ratio of BV-2:SK-ARE-*luc2* co-cultures differentially modulate neuronal NFE2L2 activity. Luciferase activity is expressed as RLU is reported on the graph as FC on monoculture; bars are mean values ±SD of n=5 measures in triplicate. \*\*\**P*<0.001 by one-way ANOVA followed by Dunnet’s multiple comparisons test.

**Supplementary Figure 4:**
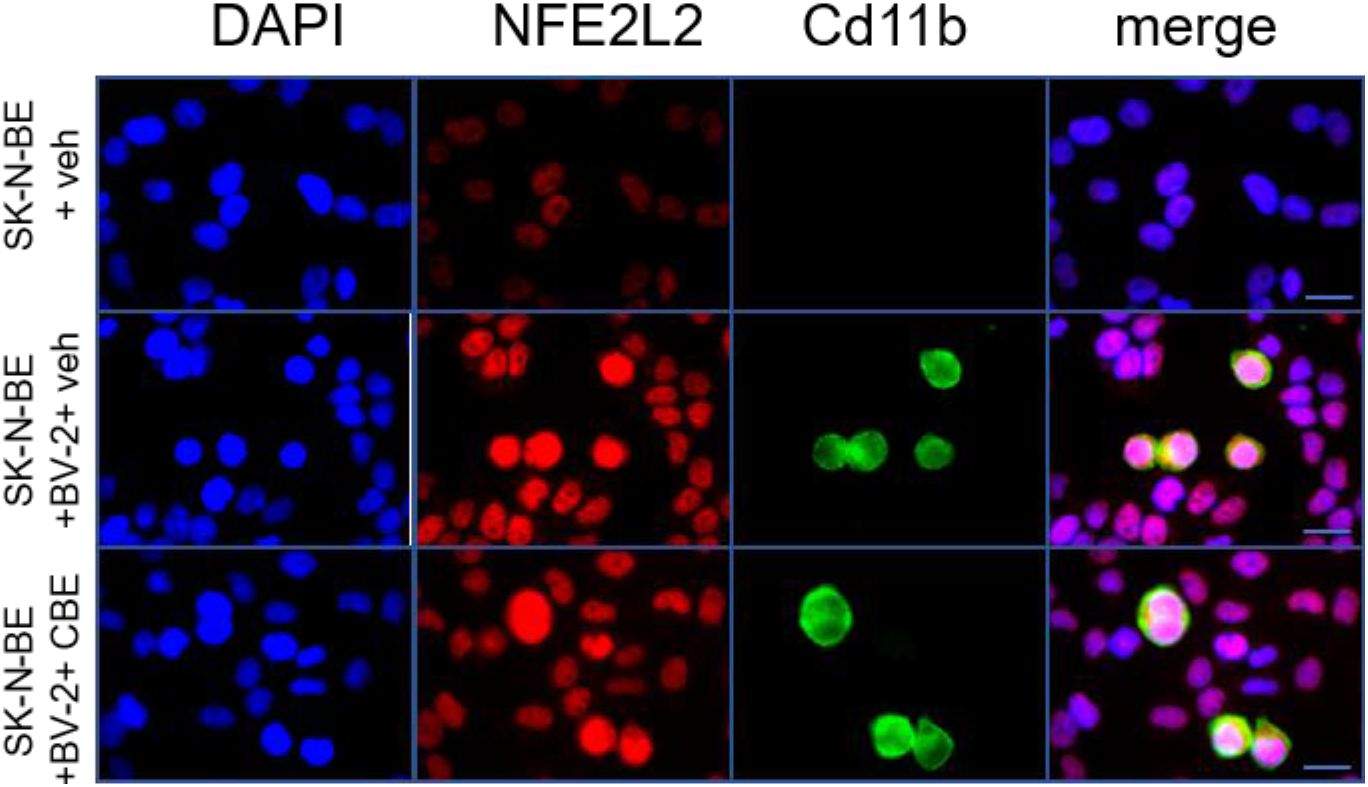
Nuclear localization of NFE2L2 in neuronal-microglia culture treated with CBE. Representative immunocytochemistry analysis of SK-N-BE and BV-2 cell lines in monoculture and co-culture, treated with 200 μM CBE or vehicle for 48 hours; cells were co-stained with anti- NFE2L2 (red) and anti-CD11b antibodies (green), and with DAPI (blue).

**Supplementary Figure 5:**
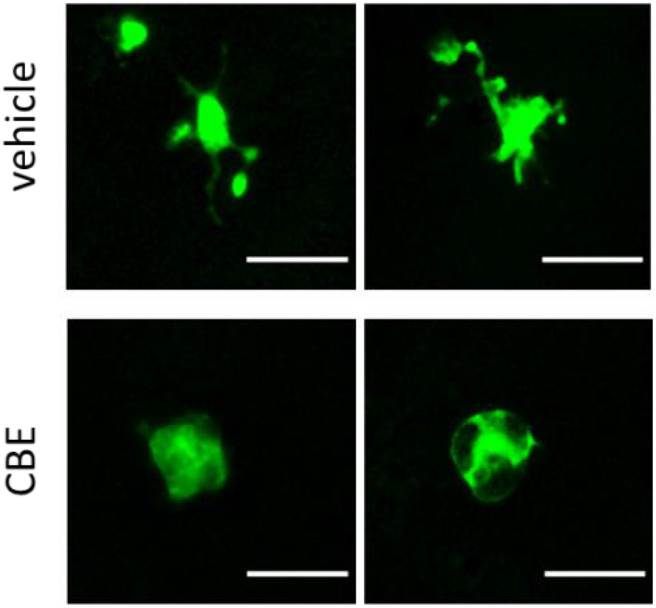
GCase inhibitions in primary microglia. Representative images of primary microglia marked with GFP showing that the treatment with 200 μM CBE for 48 hours increases a microglia sub-population characterized by a round shaped morphology.

